# Stability Transitions in Discrete Tumor-Immune Dynamics under Immune Suppression

**DOI:** 10.64898/2026.01.14.699451

**Authors:** Nara Yoon, Jacob G. Scott, Young-Bin Cho

## Abstract

Despite major advances in cancer immunotherapy, many patients still experience immune escape and disease progression. Understanding the dynamical mechanisms governing the transition between tumor control and immune escape is therefore essential for improving therapeutic strategies. Bifurcation theory and stability analysis provide a mathematical framework for explaining how gradual parameter changes can produce sudden qualitative transitions in biological systems. In the context of tumor–immune interactions, such transitions may correspond to critical thresholds separating immune-limited tumor control from immune-escape behavior.

In this study, we investigate a discrete-time model of the cancer–immunity cycle applying the local stability analysis to the system equilibria based on the trace and determinant of the Jacobian matrices. The model incorporates immune suppression through a parameter representing tumor-mediated immune evasion, including mechanisms related to immune checkpoint pathways such as PD-1/PD-L1 signaling. The system exhibits a saddle-node bifurcation associated with the appearance and disappearance of nontrivial equilibria, under weak immune conditions. In addition, under strong immune conditions, the model demonstrates a distinct stability transition in which a stable spiral equilibrium (spiral sink) loses stability and becomes an unstable spiral (spiral source), resulting in the loss of stable tumor-control dynamics before equilibrium disappearance occurs.

Additional mathematical analysis and numerical investigations indicate that the system does not generate stable non-equilibrium attractors such as limit cycles over the parameter ranges considered. Consequently, the stable equilibrium remains the only stable attractor in the model, emphasizing the importance of maintaining equilibrium stability for effective immune-mediated tumor suppression. Also, further parameter analyses reveal that both equilibrium existence and stability are highly sensitive near bifurcation boundaries, reflecting the delicate balance between tumor proliferation and immune activation.

Overall, this work provides a mathematical framework for distinguishing equilibrium existence from effective tumor control in discrete tumor–immune systems. Beyond the specific model considered, the results highlight the importance of stability analysis in understanding immune escape and may contribute to future approaches in adaptive and personalized immunotherapy modeling.

## 1 Introduction

The interaction between tumor cells and the immune system plays a fundamental role in cancer progression and therapeutic response. While the immune system is capable of recognizing and eliminating malignant cells through immune surveillance, tumors may develop mechanisms to evade immune-mediated destruction, leading to uncontrolled growth and disease progression. This balance between immune control and immune escape has therefore become a central topic in cancer biology and immunotherapy research. In particular, immune checkpoint pathways such as PD-1/PD-L1 demonstrate how tumors suppress antitumor immune activity through inhibitory signaling mechanisms, while therapeutic blockade of these pathways can partially restore immune function in some patients [1, 2, 3]. However, clinical responses remain highly heterogeneous, and many tumors eventually develop resistance or fail to respond altogether [4, 5]. These observations highlight the need for quantitative frameworks capable of describing the dynamical interaction among tumor growth, immune suppression, and therapeutic intervention, as well as supporting the development of more effective and personalize treatment strategies.

Among various quantitative approaches, discrete-time dynamical systems provide a particularly natural framework for studying biological and therapeutic processes that evolve intermittently or are administered at discrete time points. Fractionated radiotherapy, periodic immunotherapy dosing, and delayed immune activation are examples of phenomena that can be represented more naturally through difference equations than continuous-time formulations. Discrete models also permit direct analysis of iterative tumor–immune interactions and their long-term dynamical behavior. [6, 7, 8, 9] Bifurcation analysis investigates qualitative transitions in dynamical systems as parameters vary. [10, 11, 12] In biological systems, such transitions may correspond to critical thresholds separating tumor control from immune escape. Previous analyses of tumor–immune systems have focused on the appearance or disappearance of equilibria, particularly through saddle-node bifurcation mechanisms. [9] However, the existence of an equilibrium alone does not necessarily guarantee biologically achievable tumor control; stability of the equilibrium is equally essential. Changes in equilibrium stability may occur independently from changes in equilibrium existence, potentially leading to qualitatively different biological outcomes.

In this study, we investigate the bifurcation structure of a discrete-time model [12] of the cancer–immunity cycle previously proposed by Cho et al. [9]. After introducing the discrete tumor-immune model in Section 2, we analyze the existence of equilibria in Section 3 and their sability properties in Section 4 as the immune suppression parameter varies. Section 5 further explores the bifurcation behavior with respect to additional model parameters. Based on the discovered bifurcation conditions and stability transitions, and circumstance to form bifurcations, we then discuss its biological significance in the context of immunotherapy design, and conclude the research in Section 6.

## 2 Mathematical model: cancer-immunity cycle

Recent research on cancer dynamics has investigated the interaction between cancer cell population *T* and immune cell (lymphocyte) population *L* using a discrete-time dynamical system. [6, 9] A simplified version of this model, diagrammed in Figure 1, captures these essential biological mechanisms: baseline proliferation of cancer cells (*µ >* 0), decay of lymphocytes (0 *< λ <* 1), tumor-activated immune infiltration into tumor microenvironment (*ρ >* 0), and immune response *Z* = *Z*(*T, L*) in terms of lymphocytes density and the surface area of tumor. The system is formulated as follows:

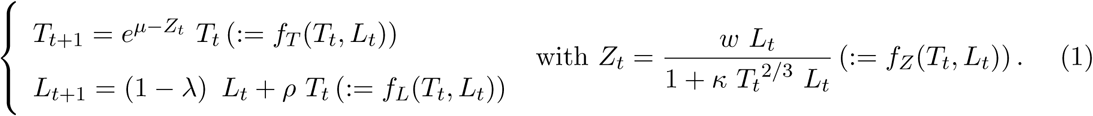

**Figure 1:**
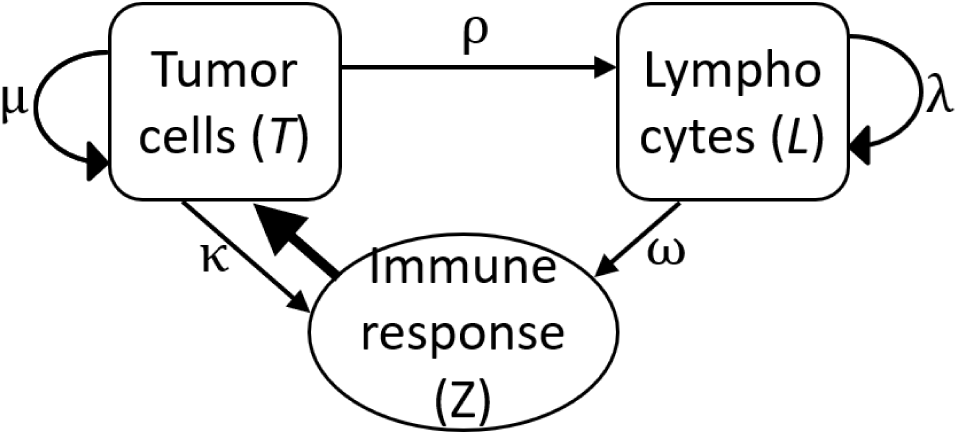
Diagram of cancer-immunity cycle. Here, *µ* and *λ* represent turnover of cancer cells and lymphocytes, respectively, *ρ* for immune cell infiltration into tumor, *ω* for immune response against cancer cells, and *κ* for immune evasion.

Specifically, in the formation of *Z*(*T, L*), we parametrized the anti-tumor immune effect (*ω >* 0), and the mechanism an opposite effect which is the immune evasion (*κ >* 0). *κ* accounts for the overall degree of immune escape exerted by the tumor microenvironment, including a range of immune suppressive mechanisms, such as immune checkpoint signaling, inhibitory cytokine activity, and other pathways that attenuate antitumor immune responses and promote tumor progression.

A prominent example is the programmed cell death protein 1 (PD-1) pathway, in which PD-1 expressed on activated immune cells binds to its ligand PD-L1 on tumor cells, suppressing immune activity. [1, 13, 2]

Therapeutic interventions targeting immune evasion mechanisms are represented in the model by decreasing the value of *κ*, thereby reducing immune suppression and restoring immune-mediated tumor control. In the next two sections, we will study the behaviors of System (1) as *κ* changes.

## 3 Emergence of non-zero equilibria with low *κ* values

Previous studies [9] have shown that System (1) may admit a stable nonzero equilibrium, interpreted as an immune-limited state, or fail to do so, leading to unchecked tumor growth or immune escape. Also, the bifurcation behavior associated with varying *κ* has been numerically studied in terms of the existence of a nonzero equilibrium. Here, we analyze it further by developing a rigorous equilibrium framework. We focus on the change in terms of the frequency of equilibria in this chapter, deferring the discussion about their stability to the next section.

System (1) admits the trivial equilibrium (0, 0), since *f* (0, 0) = 0 and *g*(0, 0) = 0 hold for any value of *κ*. Nontrivial (nonzero) equilibia (*T* ^+^*, L*^+^) exist if and only if the following conditions are satisfied:

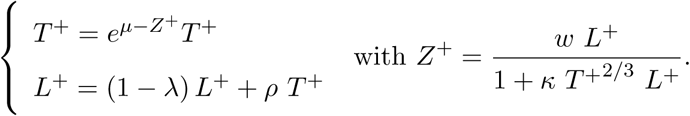

This system simplifies to

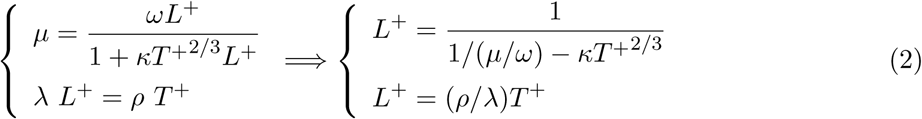

where the last step equates two expressions for *L*^+^. To facilitate the analysis of nontrivial equilibria, we define two functions *a_κ_* and *b*, implied from Equation (2):

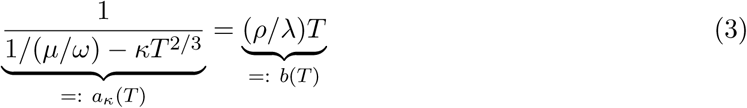

The equilibrium points correspond to the intersections of *a_κ_*(*T*) and *b*(*T*). As illustrated in Figure 2 (a), the function *a_κ_*(*T*) is concave upward (*a*^′^*_κ_*^′^(*T*) *>* 0) while *b*(*T*) is linear. The number of intersections (thus the number of nontrivial equilibria) is up to two, and depends critically on *κ*. Specifically, there exist a threshold *κ*^∗^ such that two positive equilibria exist for 0 *< κ < κ*^∗^, only one equilibrium at *κ* = 0 and at *κ* = *κ*^∗^, and no equilirium for *κ > κ*^∗^ (except the trivial equilibrium). At the critical value *κ* = *κ*^∗^, the two equilibria coalesce into a single equilibrium corresponding to the tangential intersection where the derivatives coincide, i.e., *a_κ_*∗ (*T*) = *b*(*T*) and *a*^′^*_κ_*∗ (*T*) = *b*^′^(*T*). This condition yields an explicit formula for *κ*^∗^:

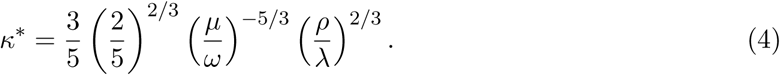

**Figure 2:**
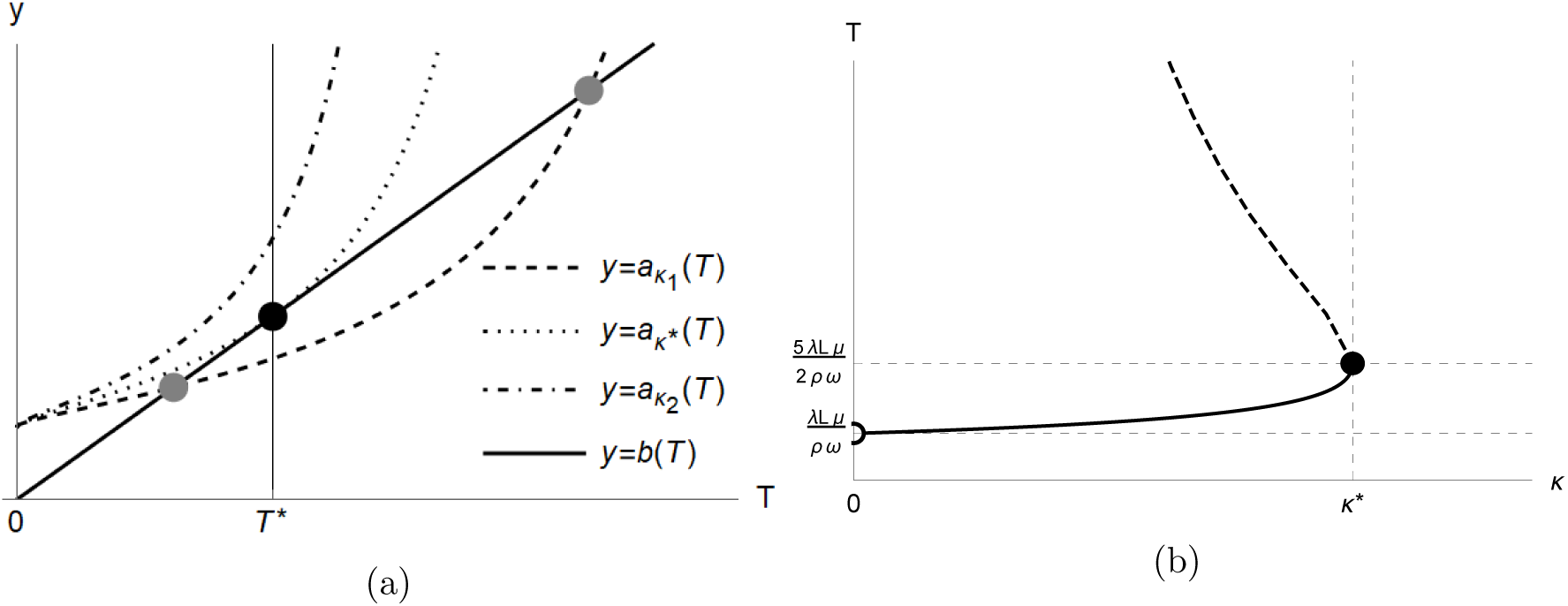
Nontrivial equilibria of System (1). (a) Curves of *a_κ_*(*T*) and *b*(*T*) (from Equation (3)) for varying *κ*. There are two intersections when *κ* = *κ*_1_ such that *κ*_1_ ∈ (0*, κ*^∗^), one intersection at *κ* = *κ*^∗^, and no intersection when *κ* = *κ*_2_ such that *κ*_2_ ∈ (*κ*^∗^, ∞). (b) Curve of nontrivial equilibrium *T* ^+^ as a function of *κ*. *T* axis is the vertical asymptote since *κ* → 0^+^ as *T* ^+^ → ∞.

In addition to when *κ* = *κ*^∗^, a single nonzero equilibrium exists at *T* ^+^ = *<u>^µ/ω^</u>* when *κ* = 0 from Equation (2). Depending on the stability of the equilibria around *κ*^∗^, a saddle-node bifurcation occurs at *κ* = *κ*^∗^, and this crucial value *κ*^∗^ is coincides with the transition moment previously identified in Cho et al. [9].

## 4 Stability transition with varying *κ*

Based on the equilibria identified in the previous section, we extend this analysis by rigorously determining the types of the equilibria, especially in terms of their stability properties, employing the bifurcation theory summarized in a diagram on Figure 3, and detailed in Appendix A. We characterize the local stability of equilibria in terms of the trace and determinant of the Jacobian matrix and investigate how changes in *κ* induce bifurcations in the cancer-immunity dynamics.

**Figure 3:**
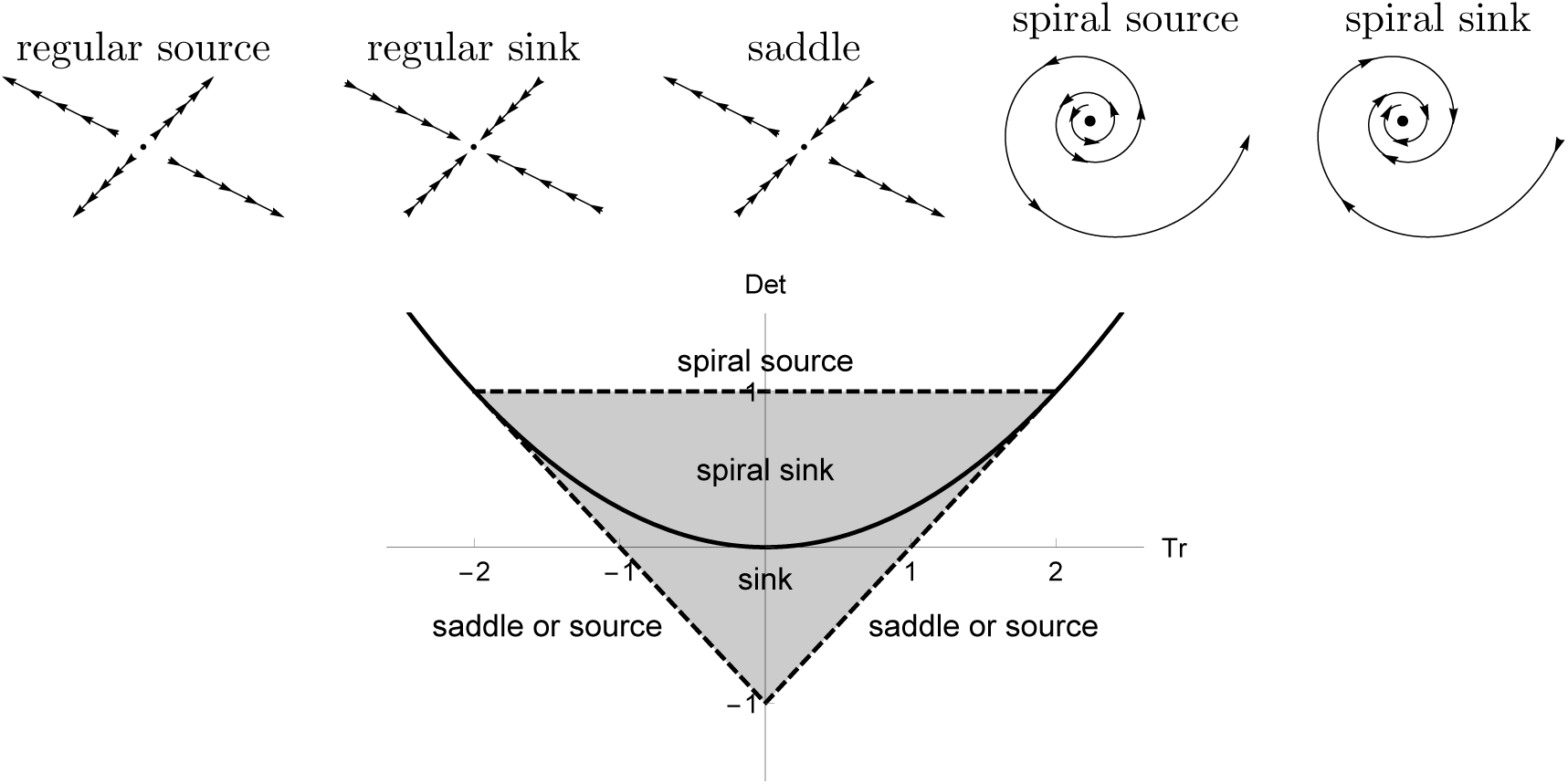
**Trace-Determinant (***Tr* − *Det***) planes** separated by the equilibrium types for two-dimensional discrete systems (9). The separating boundaries are *Tr*^2^ − 4*Det* = 0, *Det* = |*T r*| − 1 and *Det* = 1. The shaded area is are the regions of stable equilibria (sink or spiral sink).

To apply it to this work, we begin by evaluating the Jacobian matrix of System (1):

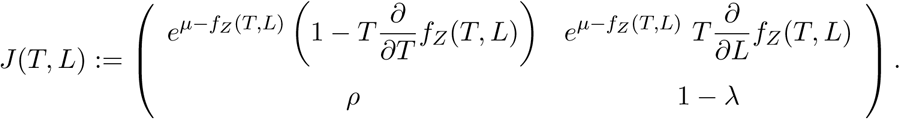

Let *Tr* and *Det* denote the trace and the determinant of *J* :

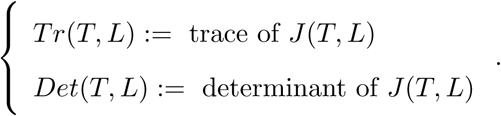

At the trivial equilibrium point (0, 0), we have

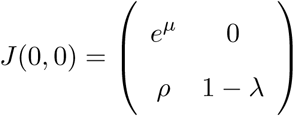

The corresponding eigenvalues are *e^µ^ >* 1 and 1 − *λ* ∈ (0, 1), implying that the origin is a saddle point and hence unstable for any value of *κ* (by Table 2 in Appendix A).

At the critical point *κ* = *κ*^∗^ (Equation (4)) from the last section, the system admits a nontrivial equilibrium at

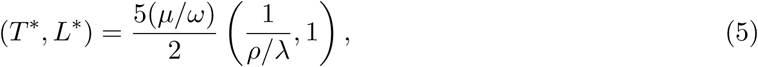

and the Jacobian evaluated at this point becomes:

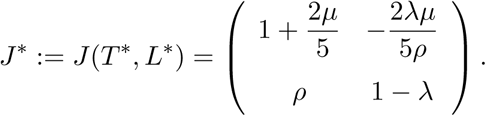

The determinant of *J* ^∗^ can be calculated as:

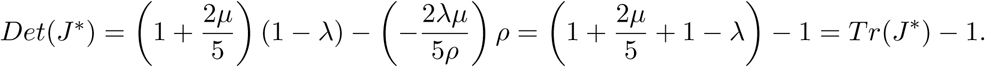

This indicates that the point (*Tr, Det*) at *κ* = *κ*^∗^ lies exactly on the line *Det* = *Tr* − 1 in the (*Tr, Det*) plane. If the trace value lies between 0 and 2 (equivalently if the determinant value lies between −1 and 1), the equilibrium lies on the boundary of the stable region represented by the inverted triangular domain in the *Tr* − *Det* diagram (Figure 3. On the top of that, if the lower- and higher-equilibrium branches extend respectively into and out of the stable region (or vice versa), the system exhibits a saddle-node bifurcation at (*T* ^∗^*, L*^∗^) (with *κ* being *κ*^∗^). An example of this bifurcation is observed when we use the parameter values of *µ* = 0.217 [14], *ρ* = 0.1, *ω* = 0.05 and *λ* = 0.335 [15]. (Figure 4) Here, *κ*^∗^ = 0.0126, and *T* ^∗^ = 36.35. For 0 *<κ < κ*^∗^, a stable non-trivial equilibrium coexists with an unstable one; for *κ > κ*^∗^, no non-trivial equilibrium exists.

**Figure 4:**
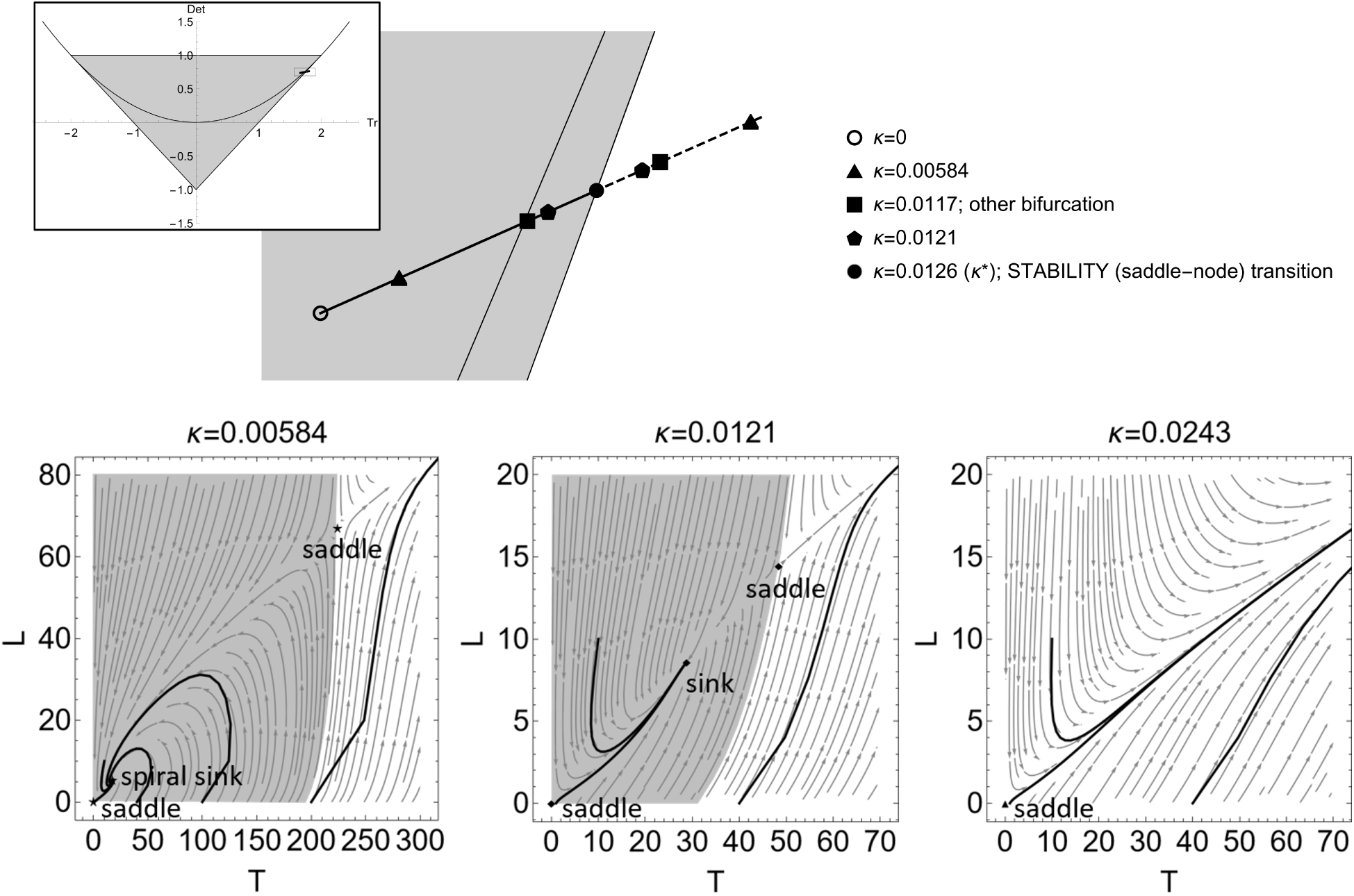
**Bifurcation diagram for a weak immune system as immune suppression (***κ***) varies,** with the parameter values from Table 1. The upper panel shows the trajectory of the nonzero equilibrium tumor size *T* ^+^ as functions of *κ* ∈ [0*, κ*^∗^] (with *κ*^∗^ = 0.0126), mapped onto the trace-determinant (*Tr* − *Det*) plane. The point corresponding to the critical bifurcation value *κ*^∗^ is indicated by a filled circle. The solid and dotted curves represent the lower (stable) and upper (unstable) equilibrium branches, respectively. Another significant shift occurs at *κ* = 0.0117, where the stable equilibrium transitions from a spiral sink to a regular sink. The bottom three panels display the direction fields of the {*T, L*} dynamics at three representative values of *κ*, selected between bifurcation points to highlight qualitatively distinct behaviors. Each panel includes trajectories from three different initial conditions. When *κ* is below the stability transition threshold (*κ*^∗^), a stable equilibrium exists, along with a region of initial conditions (shaded area) from which the tumor state converges to this equilibrium over time.

**Table 1:**
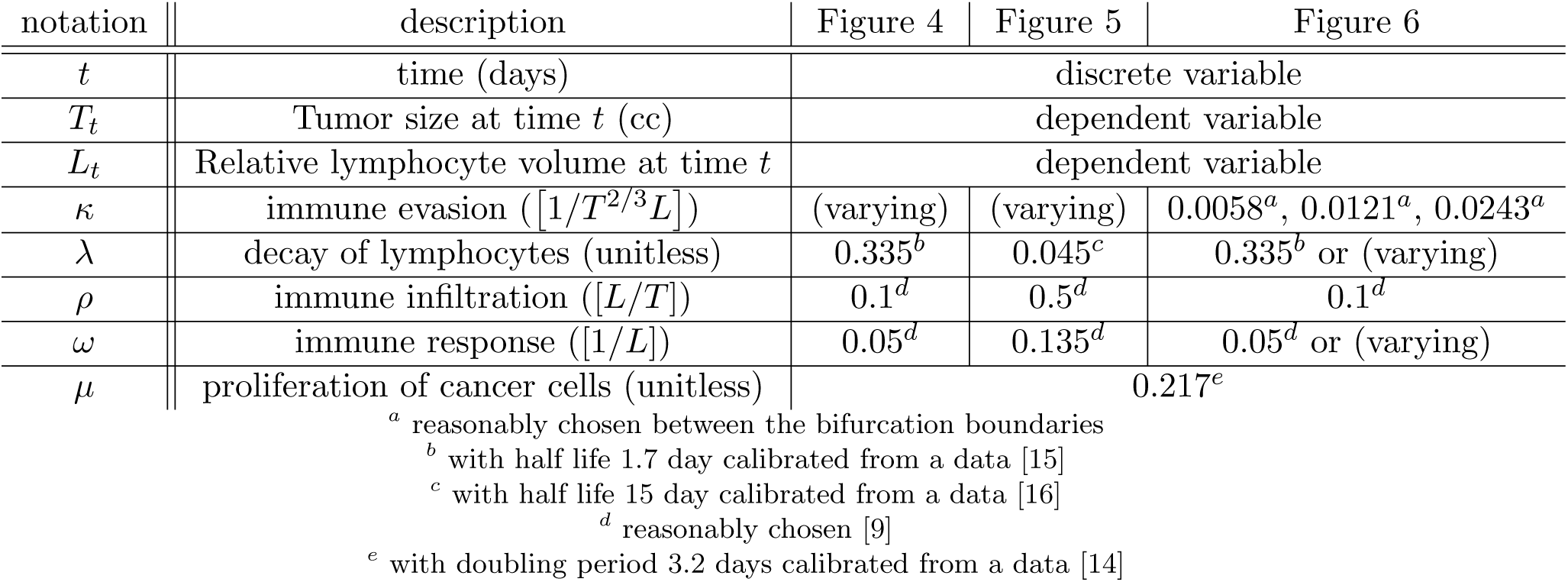
Variables and parameters in the System 1

| notation | description | Figure 4 | Figure 5 | Figure 6 |
| --- | --- | --- | --- | --- |
| $t$ | time (days) | discrete variable | | |
| $T_t$ | Tumor size at time $t$ (cc) | dependent variable | | |
| $L_t$ | Relative lymphocyte volume at time $t$ | dependent variable | | |
| $\kappa$ | immune evasion ($[1/T^{2/3}L]$) | (varying) | (varying) | $0.0058^a, 0.0121^a, 0.0243^a$ |
| $\lambda$ | decay of lymphocytes (unitless) | $0.335^b$ | $0.045^c$ | $0.335^b$ or (varying) |
| $\rho$ | immune infiltration ($[L/T]$) | $0.1^d$ | $0.5^d$ | $0.1^d$ |
| $\omega$ | immune response ($[1/L]$) | $0.05^d$ | $0.135^d$ | $0.05^d$ or (varying) |
| $\mu$ | proliferation of cancer cells (unitless) | $0.217^e$ | | |

<sup>a</sup> reasonably chosen between the bifurcation boundaries<sup>b</sup> with half life 1.7 day calibrated from a data [15]<sup>c</sup> with half life 15 day calibrated from a data [16]<sup>d</sup> reasonably chosen [9]<sup>e</sup> with doubling period 3.2 days calibrated from a data [14]

**Table 2:**
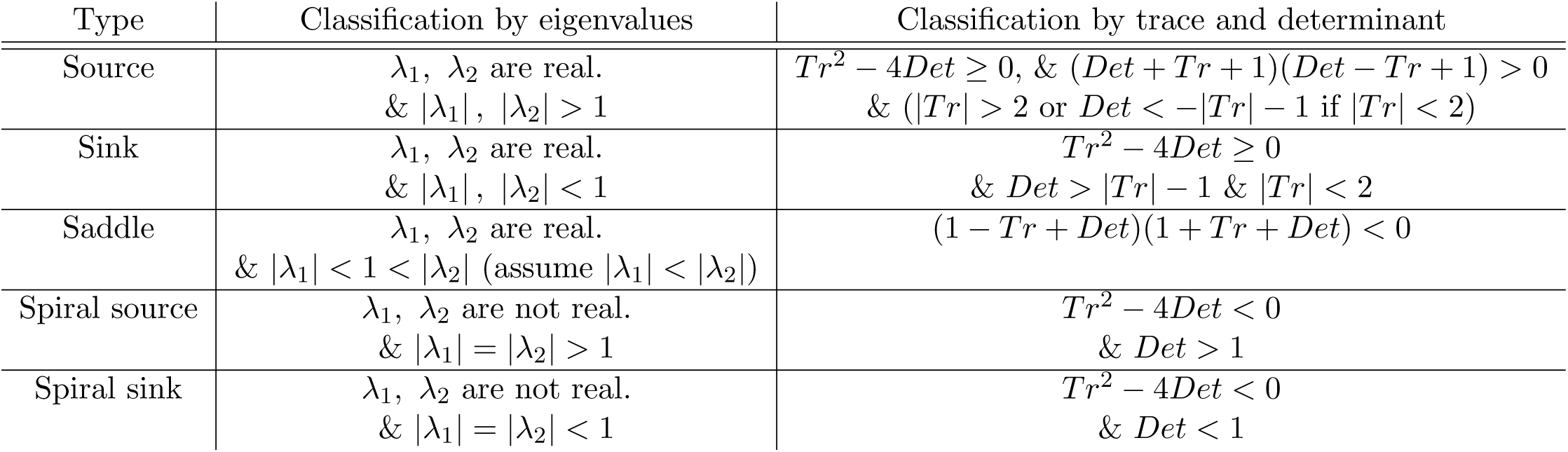
Eigenvalue analysis of equilibrium types for System (9) comprised by two difference equations with two dynamical variables. *λ*_1_ and *λ*_2_ are the eigenvalues of the Jacobian matrix at the system equilibrium, and *Tr* and *Det* are trace and determinant of the matrix.

A simulation in a stronger immune environment shows differently interesting bifurcation patterns, in that the equilibrium trajectory enters the stability region at *κ* = *κ*^∗∗^ which is lower than *κ*^∗^. The explicit expression of *κ*^∗∗^ and the corresponding value of tumor equilibrium *T* ^+^ = *T* ^∗∗^ are:

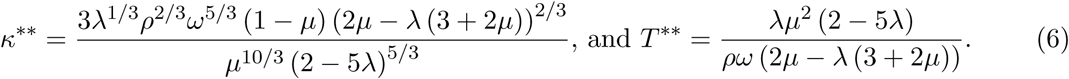

An example is shown on Figure 5. With the parameter values of *µ* = 0.217 (same with the previous example), *ρ* = 0.5 (enhanced immune infiltration), *ω* = 0.135 (increased immune-induced tumor cell killing) and *λ* = 0.045 (slower lymphocyte decay), (*Tr, Det*) trajectory corresponding to (*κ, T* ^+^) crosses the inverse triangle region of the *Tr*−*Det* diagram through the upper base (*Det* = 1) at *κ*^∗∗^ = 0.500 with *T* ^∗∗^ = 0.199 (and after then it intersects *Det* = *Tr* − 1 at *κ*^∗^ = 0.735). This shows that, if *κ*^∗∗^ *< κ < κ*^∗^, there are two nontrivial equilibria, but both are unstable. Therefore, a stable equilibrium occurs for *κ < κ*^∗∗^ ≪ *κ*^∗^ for this parameter set. This *κ*^∗∗^ is still much higher than *κ*^∗^ = 0.0126 of the weaker immune system case. However, this observation indicates that immuno-therapeutic intervention must aim to reduce immune suppression below the lower threshold to achieve a realistic possibility of cancer control. Relying soley on the existance of nonzero equilibrium, as a treatment guideline, may overestimate the effectiveness of immunotherapy, and consequently lead to unsuccessful outcome.

**Figure 5:**
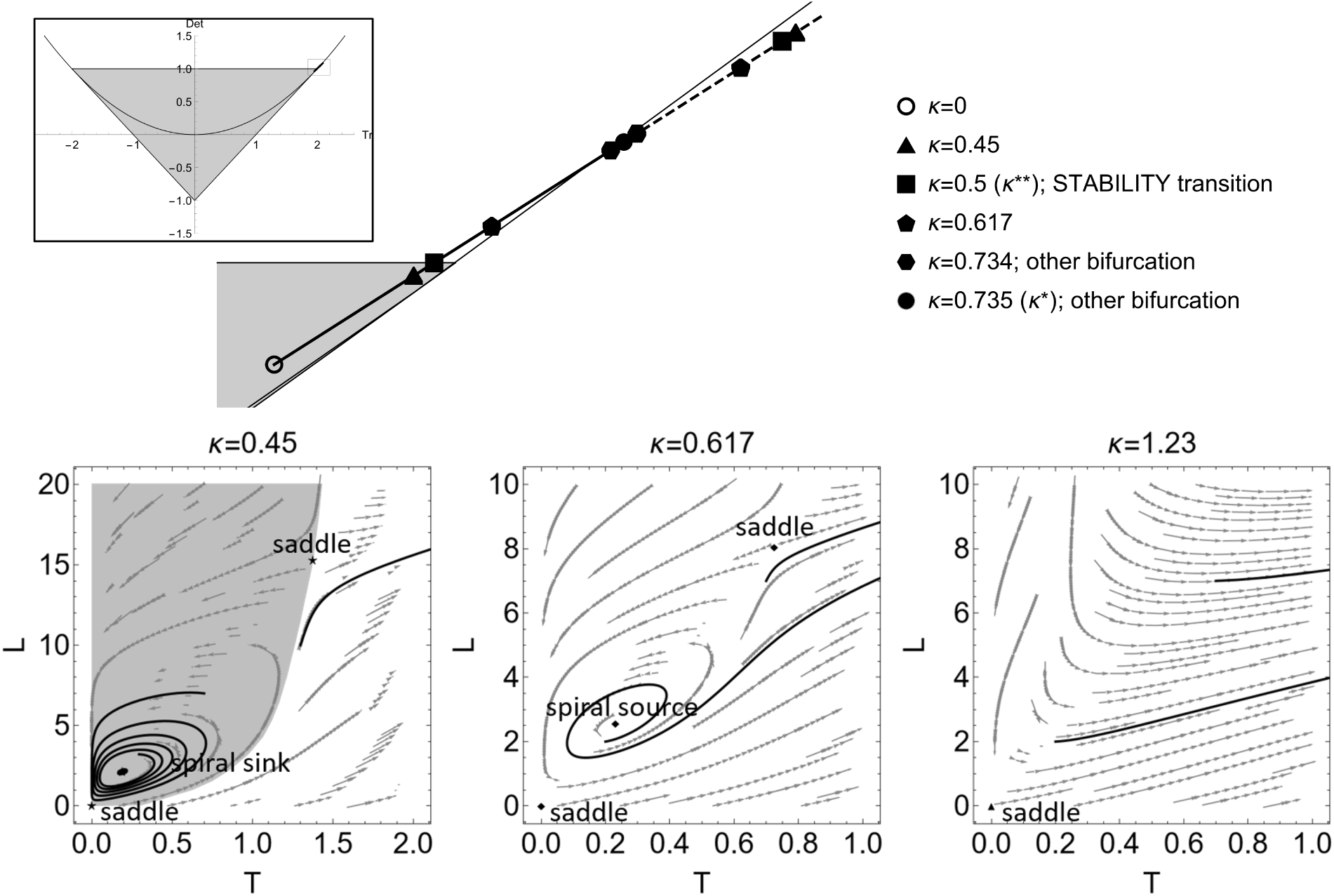
**Bifurcation diagram for a strong immune system as immune suppression (***κ***) varies,** with the parameter values from Table 1. The upper panel shows the trajectories of nonzero equilibrium tumor size *T* ^∗^(s) as *κ* ∈ [0*, κ*^∗^] (with *κ*^∗^ = 0.735), mapped onto the trace-determinant (*Tr* − *Det*) plane. The point corresponding to the bifurcation by the number of equilibrium *κ*^∗^ is indicated by a filled circle. The solid and dotted curves represent the lower equilibrium (stable or unstable) and upper equilibrium (unstable always) branches, respecrively. The stability transition occurs with *κ*^∗∗^ = 0.5 at which the lower equilibrium changes from spiral sink (stable) to spiral source (unstable). One additional transition is observed at *κ* = 0.734 where the lower (unstable) equilibrium change the type from spiral source to regular source/saddle. The bottom three panels display the direction fields of the {*T, L*} dynamics at three representative values of *κ*, selected between the transition points to highlight qualitatively distinct behaviors. Each panel includes trajectories from three different initial conditions. When *κ* is below the stability transition threshold (*κ*^∗∗^ = 0.5), a stable equilibrium exists, along with a region of initial conditions (shaded area) from which the tumor state converges to this equilibrium over time.

Additional mathematical analysis indicates that this stability transition at *κ* = *κ*^∗∗^ with *Tr*^∗∗^=1 occurs as the trajectory crosses the boundary between the stable (spiral sink) and unstable (spiral source) regions (as shown on Figure 7 in Appendix A). We want to note here that the system still does not undergo a Neimark–Sacker bifurcation which leads to the emergence of a stable limit cycle around the destabilized equilibrium. It is because the nonlinear conditions required for the emergence of an invariant closed curve are not satisfied. In the present model, nonlinear effects arise exclusively through the immune response term *Z*(*T, L*), while the remaining components are linear. Numerical investigations indicate that the overall nonlinear structure of the system is insufficient to generate sustained oscillatory dynamics or stable limit cycles.

Consequently, the stable equilibrium remains the only stable attractor observed in the system, which means that only when an effective immunotherapy reduces *κ* below *κ*^∗^ in case of weak immune systems, and below *κ*^∗∗^ in case of strong immune systems, there is a chance to control tumor growth, with the equilibrium tumor size bounded above by *T* ^∗^ and *T* ^∗∗^, respectively.

Also, one additional condition for successful immune control along with both low *κ* is that the initial tumor size is low enough to be in the convergence basin, as the multiple trajectories of {*T, L*} on Figures 4 and 5 show. The region of convergence is numerically generated, and drawn by the gray shade. An additional requirement for effective immune control, beyond the condition *κ < κ*^∗^, is that the tumor burden must remain sufficiently small. As shown in the lower panels of Figure 4, trajectories starting outside the gray region–manually found based on the numerical simulation –diverge, indicating uncontrolled tumor growth.

## 5 Sensitivity and bifurcation of other parameters

In addition to the immune suppression parameter *κ* examined in the earlier sections, our dynamical system is governed by four additional parameters : *µ*, *ω*, *ρ*, and *λ*. As demonstrated in the previous analysis (Equations (2),(3),(4),(5)), these parameters naturally form two coupled pairs: *µ* and *ω*, and *ρ* and *λ*. The ratios formed by these pairs (specifically, *µ/ω* and *ρ/λ*) play a central role in determining the qualitative dynamics of the system.

The ratio *µ/ω* can be interpreted as the net tumor proliferation rate under the influence of immune-mediated suppression, quantifying the balance between tumor growth and immune inhibition. In parallel, the ratio *ρ/λ* reflects the net rate of lymphocyte production induced by the presence of tumor cells, relative to their natural decay. Together, these ratios capture the interplay between tumor expansion and the host immune response.

To investigate the sensitivity of the systems to these critical ratios, we performed a bifurcation analysis in which one parameter in each pair (*µ* and *ρ*) was kept constant, while the corresponding partner parameter (*ω* and *λ*) was varied. For each scenario, we identify the associated bifurcation point by solving Equation (3). The resulting bifurcations are again of the saddle-node type, as illustrated on Figure 6 (a,b). Similar to the bifurcation structure depicted in Figure 2 (b), the bifurcation point is on the line of *Det* = *Tr* − 1, with the solution branches remaining close to the separatrix boundaries of equilibrium types on the trace-determinant plane.

**Figure 6:**
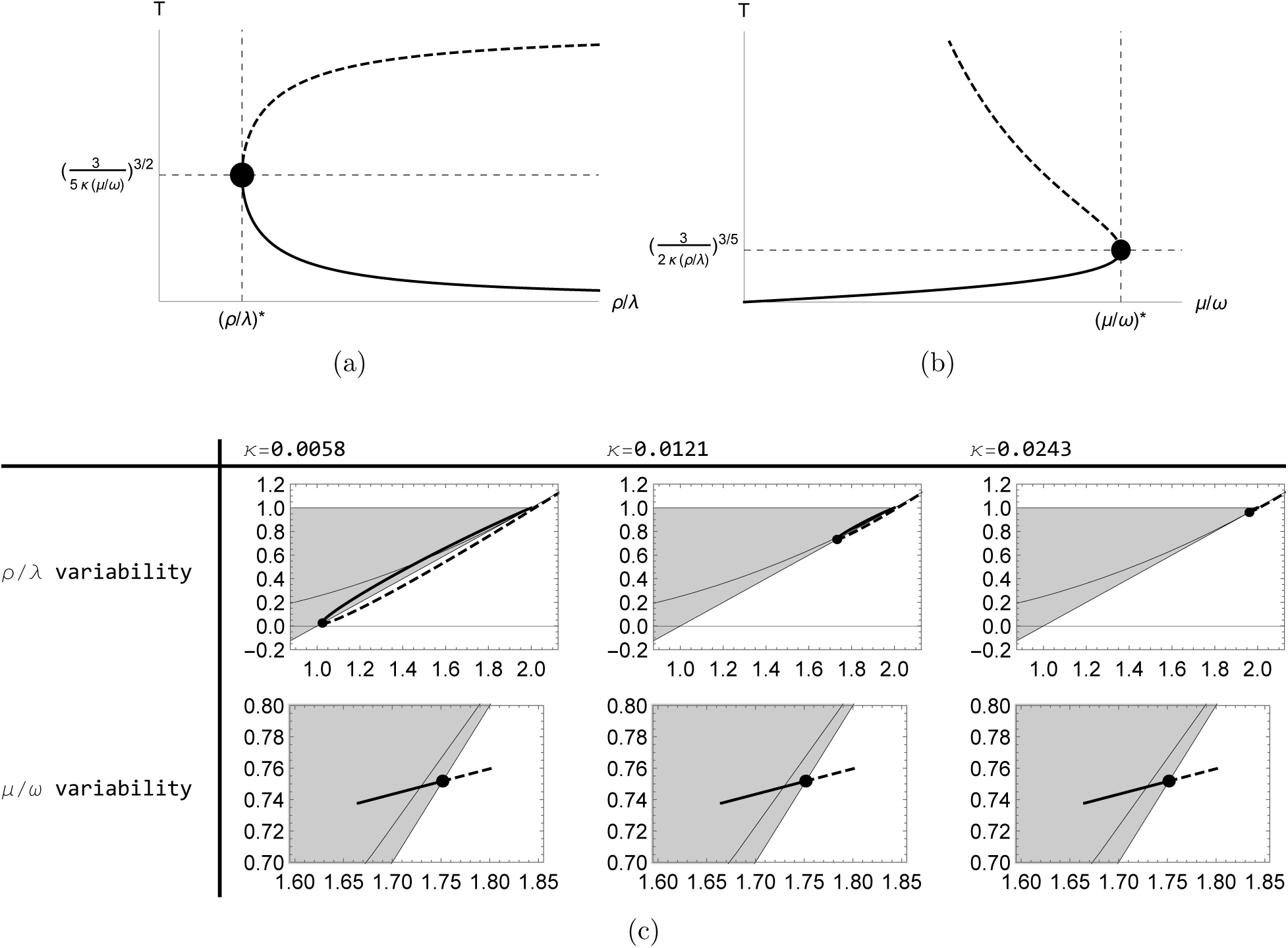
Saddle-node bifurcation as. *ρ/λ* **and** *µ/ω* **varies.** Panel (a) and Panel (b) shows the curve nonzero equilibria for the two parameter set. Panel (c) shows the equilibrium trajectories transferred onto the trace-determinant plane with three different *κ* values. In the analysis of *ρ/λ*, only *λ* is varying with *ρ* (and all the other parameters) being fixed, and in the study of *µ/ω* variability, only *ω* is varying with *µ* (and all the other parameters) being fixed. Except the parameters to be explored, the other parameter values are same with the exercise of the weak immune system (Figure 4).

**Figure 7:**
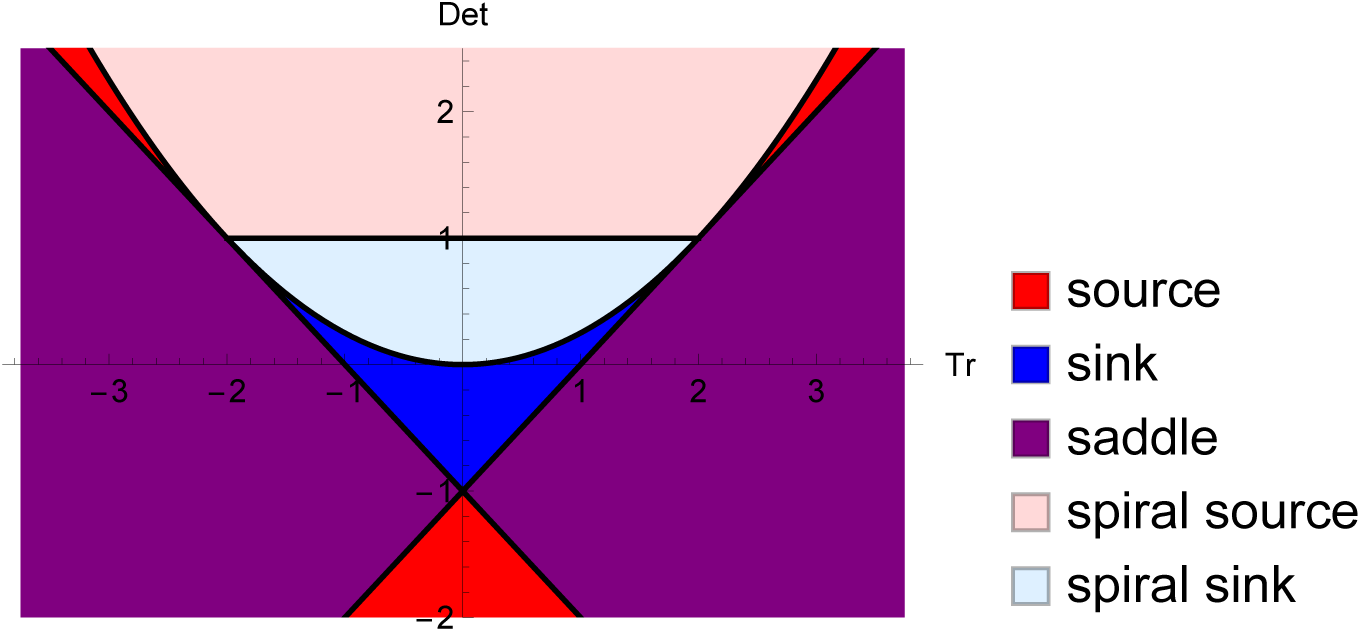
**Trace-Determinant (***Tr* − *Det***) plane** of two-dimensional discrete system, separated by the equilibrium type. The type of an equilibrium is classified by the trace and determinant values of the Jacobian matrix of the system evaluated at an equilibrium. Above the parabola (*Tr*^2^ − 4*Det* = 0) is the region relevant to spiral source/sink, below it is relevant to regular sink, source or saddle.

Interestingly, our analysis reveals distinct behaviors for the two ratios. *ρ/λ* generates an equilibrium trajectory that extends along the bifurcation boundary in the *Tr* − *Det* plane. (Figure 6 (c)) In contrast, variations in *µ/ω* drive the trajectory nearly perpendicular to the bifurcation boundary, indicating that changes in this ratio exert a stronger influence on the system’s qualitative state. From a practical standpoint, this implies that tumor proliferation dynamics, as governed by *µ/ω* constitute a more critical control parameter for inducing qualitative transitions in the system than the lymphocyte turnover ratio *ρ/λ*. This result aligns with biological intuition: tumor proliferation serves as the primary driver of the cancer-immune system interaction, where as the immune response operates as a downstream mechanism attempting to restrain tumor growth.

Taken together, these findings highlight the need to account for the balance between proliferation and immune activity in mathematical models of tumor-immune dynamics, with direct implications for designing targeted interventions that shift the system toward disease control.

## 6 Discussion and conclusions

In this study, we analyzed the bifurcation structure of a discrete-time model of the can-cer–immunity cycle under immune suppression. By combining equilibrium analysis with local stability classification on the trace–determinant plane, we investigated the dynamical transitions between immune-limited and immune-escape tumor states. Our results demonstrate that the qualitative behavior of the system is characterized through both the existence of equilibria and their stability transitions, which play an equally important role in determining whether effective immune-mediated tumor control can be maintained.

The model exhibits saddle-node bifurcations associated with the appearance and disappearance of nontrivial equilibria, under weak immune conditions. The loss of equilibrium stability occurs simultaneously with equilibrium annihilation, producing an abrupt transition from tumor control to immune escape. In contrast, under strong immune conditions, the stable equilibrium may lose stability before disapperance of nontrivial equilibrium. This earlier stability transition implies that the existence of a tumor-control equilibrium does not necessarily guarantee biologically achievable tumor suppression. Consequently, equilibrium-based criteria alone may overestimate the effectiveness of therapeutic strategies aimed at reducing immune suppression.

Additionally, although the observed stability transition under strong immune conditions involves eigenvalues crossing from the spiral sink region into the spiral source region of the trace–determinant plane, numerical investigations over the parameter ranges considered did not reveal sustained oscillatory dynamics or stable non-equilibrium attractors. The nonlinear structure of the present model appears insufficient to generate invariant closed curves or persistent oscillatory tumor behavior. Consequently, the stable equilibrium remains the only stable attractor identified in the parameter regimes studied.

Biologically, the immune suppression parameter *κ* represents the overall strength of tumor-mediated immune evasion, including mechanisms such as immune checkpoint signaling and suppressive interactions within the tumor microenvironment. In this interpretation, reducing *κ* corresponds to therapies that restore antitumor immune activity, including immune checkpoint blockade. Our analysis suggests that successful immune restoration requires not only the existence of a favorable equilibrium state, but also preservation of its dynamical stability. This observation may help explain heterogeneous responses to immunotherapy, in which some patients fail to achieve durable tumor control despite partial immune activation.

Importantly, however, even when *κ* is lower than the bifurcation point, effective immune control depends on initial conditions; large tumor burdens may lie outside the attraction basin of stable equilibrium, resulting in unbounded growth. Therefore, in addition to the therapy controlling immune evasion, regulating tumor size (by surgery, radiation, chemotherapy, etc.) is required for the success of cancer therapy.

Additional sensitivity analyses further revealed that the system dynamics are strongly influenced by the balance between tumor proliferation and immune activation. In particular, the relative magnitudes of tumor growth and lymphocyte recruitment parameters significantly affect both equilibrium existence and stability. These results emphasize that tumor–immune dynamics remain highly sensitive near bifurcation boundaries, where small parameter changes may produce substantial qualitative differences in long-term behavior.

By the trace-determinant bifurcation framework, this cancer therapy research provides a mathematical toolset for more accurate prediction of treatment outcomes. The bifurcation structures revealed here offer a mechanistic interpretation of immune escape and support a rational basis for adjusting therapeutic parameters to sustain immune-mediated tumor control. We acknowledge the clinical limitation that immunotherapies are typically administered for finite durations due to immune-related adverse events, cost, or diminishing efficacy over time [5, 3, 4]. Consequently, our work can be applicable in personalized medicine by applying it to the patients with mild immune

suppression, so that a reasonable amount of medicine can change the game. Also, consequently, this emphasizes the importance of combining immunotherapy with tumor-debulking strategies such as surgery or radiotherapy, which may shift system dynamics toward favorable control regions [17]. While this modeling and analysis provide valuable insights into cancer-immune dynamics and potential clinical applications, several limitations should be acknowledged, some of which offer directions for future research. First, the bifurcation theory applied to discrete systems relies on local linear approximations (as shown in Equation (7)), which may be inaccurate near bifurcation boundaries or separatrices that define convergence basins. [11, 12] Consequently, global nonlinear behaviors could deviate from those predicted by local analysis.

Moreover, the model employs several simplifications, including aggregate immune populations, static parameters, and the absence of spatial or stochastic features, that limit its direct translational relevance. These simplifications are common in mathematical oncology to enable analytic tractability [18, 7], though they omit critical factors such as immune cell heterogeneity and tumor microenvironment complexity. Nonetheless, the structural insights gained from this reduced model can serve as a foundation for future extensions incorporating patient-specific data and biologically realistic features.

Future work may include dynamic parameterization, spatial modeling, and differentiation between immune cell subtypes. Spatial extensions, for example, can capture immune infiltration patterns and localized immunosuppression [19]. Additionally, we aim to integrate the pharmacodynamics of immune checkpoint inhibitors into the model, allowing the simulation of dosing schedules, therapy durations, and treatment discontinuation effects [20, 17]. Such developments could enhance the predictive and prescriptive power of bifurcation-based analysis, supporting optimized and personalized immunotherapy strategies.

### Declaration of generative AI and AI-assisted technologies in the manuscript preparation process

During the preparation of this work the authors used ChatGPT based on GPT-5.2 in order to refine the language and structural coherence. After using this tool/service, the authors reviewed and edited the content as needed and take full responsibility for the content of the published article.

## Appendix A Overview of stability of discrete systems

One of the standard discrete systems describes the present state based on the state in one time step backward. Let us represent the general form of the system for *n* state variables *X_t_* = 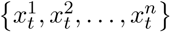 in a system of difference equations with the update function *F* : R*^n^* → R*^n^*:

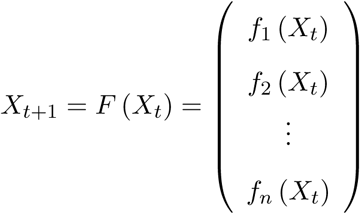

for evenly spaced distinct time points *t* ∈ {0, 1, 2, 3*, …*}. To analyze the local stability of the system equilibrium *X_eq_*(such that *X_eq_*= *F* (*X_eq_*)), let us denote Δ*X_t_* := *X_t_* − *X*_eq_ that represents the deviation from equilibrium. Then, using a first-order Taylor expansion near *X*_eq_ (i.e., *X_t_* ≈ *X*_eq_ and |Δ*X_t_*| ≈ 0), we obtain:

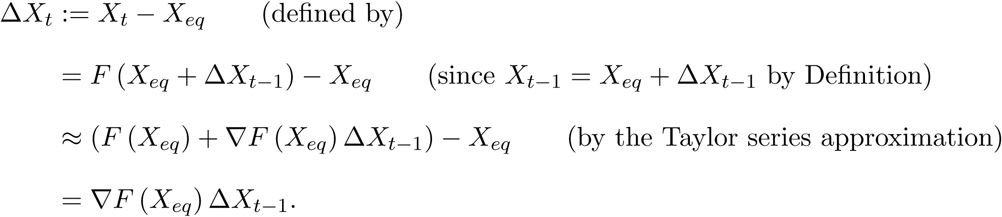

Iterating this relation yields the approximation

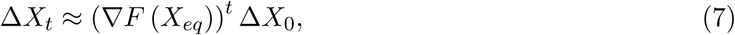

And

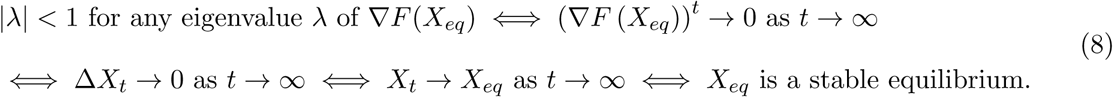

Therefore, *X_eq_* is locally asymptotically stable if and only if all eigenvalues of ∇*F* (*X_eq_*) lie strictly within the unit circle in the unit *n*-sphere (i.e., have modulus less than 1).

### A.1 Trace-determinant bifurcation plane for two dimensional discrete systems

For two dimensional real system *F* : R^2^ → R^2^, the classification criterion of equilibrium types based on the eigenvalues of ∇*F* (*X_eq_*), from the previous section, can be transformed into the criterion based on the trace *Tr* and determinant *Det* of ∇*F* (*X_eq_*). Since both *Tr* and *Det* are real always, {*Tr, Det*}-based method is more practical, in that we can visualize all the sections of equilibrium types on two dimensional Cartesian coordination system. (Figure 3 (b))

Let us start with the general form of a system of two difference equations (*f* and *g*) for two dynamic variables (*x* and *y*):

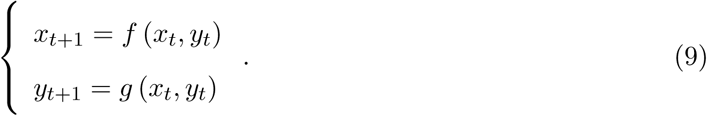

Its Jacobian matrix is:

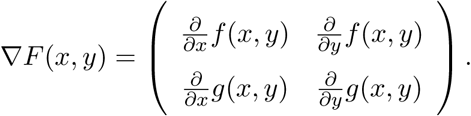

The eigenvalues of ∇*F*, *λ*_1,2_, are the root(s) of the characteristic polynomial,

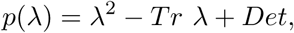

where 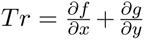 and 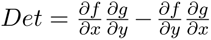 are the trace and the determinant of ∇*F*, respectively.

Based on these two measures, we derive two classification standards.

- Standard 1: Exponential behavior with real *λ*_1_,_2_ vs. oscillatory spiral behavior with non-real *λ*_1_,_2_.
- Standard 2: Stability with |*λ*_1_,_2_| *<* 1 vs. instability with |*λ*_1_| *>* 1 or |*λ*_2_| *>* 1. (by Criterion (8))

Combining the two standards, the way to classify the equilibrium types is summarized in Table 2, and is visualized in *Tr* − *Det* plane in Figure 3. The two types of exponentially unstable equilibrium, source and saddle, are not distinguished in Figure 3, due to the complexity of their analytic differences more than necessary. Instead, thorough mathematical analysis of the equilibrium classification is described in Appendix A.

### A.2 Saddle and Source areas on Trace-Determinant plane

In Chapter 2, we clarified the area of the trace-determinant plane relevant to each equilibrium type of exponential sink, exponential source/saddle, spiral sink, and spiral source. In this document, we elaborate the details of the mathematical analysis on the separation of the areas relevant to the equilibrium types.

Let us revisit the characteristic equation with the trace and determinant of the Jacobian matrix:

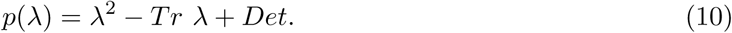

Since the zeroes of *p*(*λ*) are equal to the eigen values of the Jacobian, we can derive the following equivalences:

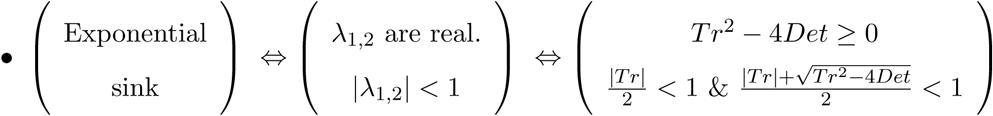

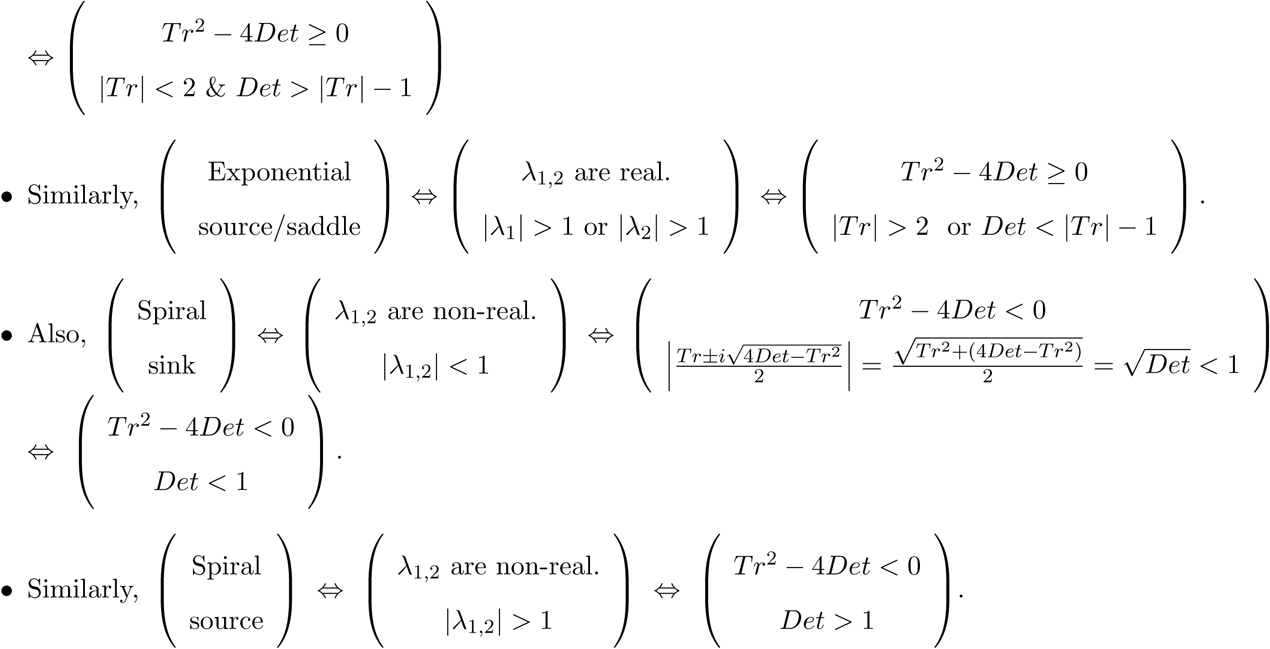

((1 − *Tr* + *Det*)(1 + *Tr* + *Det*) *<* 0), which means that the rest of the region of “Exponential source/saddle” is for “Exponential source”.

On the top of the separations in terms of equilibrium type from Figure 3, Figure 7 shows the area of saddle and source equilibria separately. The area of the exponential source is easily clarified by dropping the area for saddle type from the combined area of both nbnn types.

The following are detailed graphs of the two case studies of tumor bifurcation under the two different immune environment (shown earlier on Figures 4 and 5). Beside the stability transition, in the case of the weak immune system (Figure 8), there is an additional bifurcation as the stable equilibrium crossing the borderline of sink region and spiral sink region. In the case of the strong immune system (Figure 8), in addition to the stability transition and the disappearance of nontrivial equilibrium, there is an additional bifurcations occurring as an equilibrium remains unstable but changes the type from spiral source to source.

**Figure 8:**
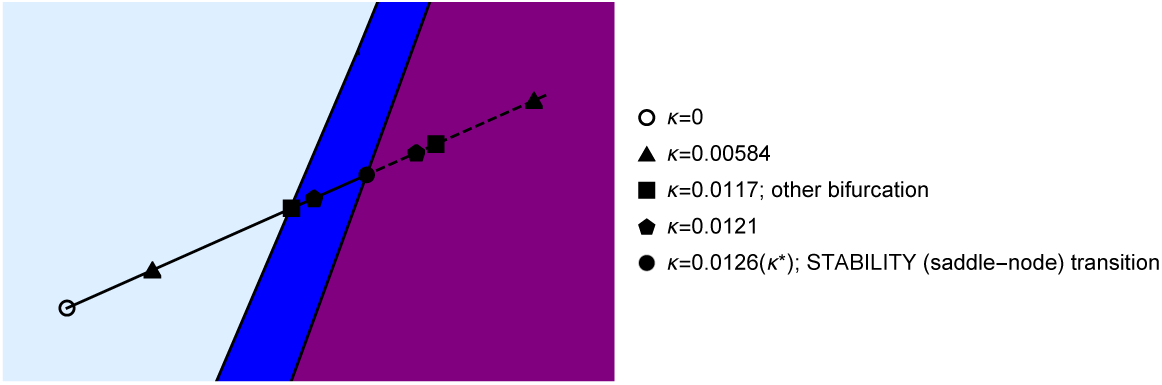
Bifurcation diagram for the weak immune system (Figure 4) on the detailed *Tr* − *Det* sections.

**Figure 9:**
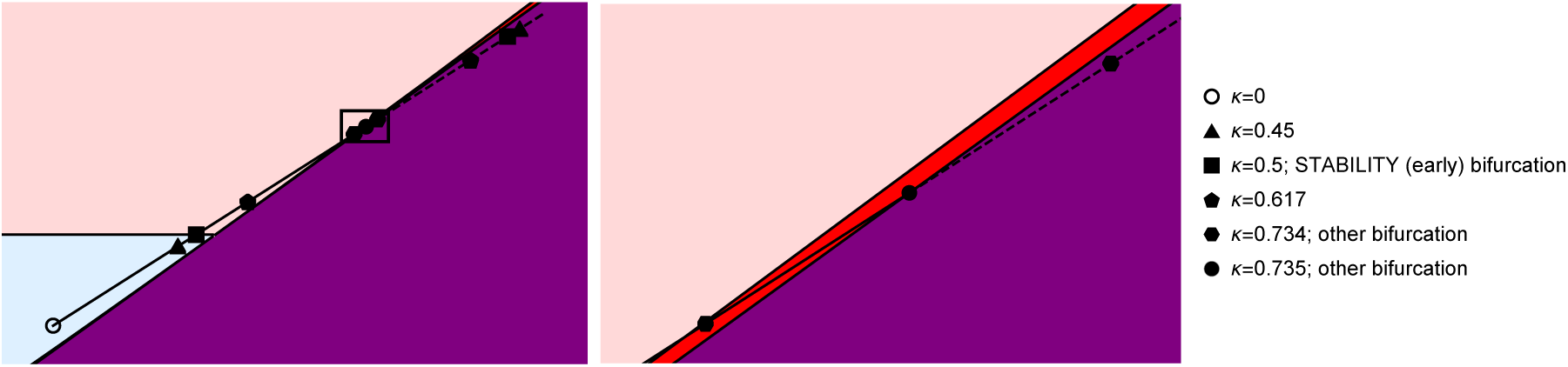
Bifurcation diagram for the strong immune system on the detailed. *Tr* − *Det* **sections.** The left panel is on the same axes range with Figure 5, and the right one is on a narrower range equivalent to the small box on the left panel manifesting the “other” bifurcation point passing a bifurcation boundary.

### Appendix B Programming code

All simulations and figures were generated using Wolfram Mathematica version 12.1. The computational codes and scripts used in this study are publicly available at: https://github.com/nryoon12/cancer-immune-cycle.

